# Senescent cells and macrophages cooperate through a multi-kinase signaling network to promote intestinal transformation in Drosophila

**DOI:** 10.1101/2023.05.15.540869

**Authors:** Ishwaree Datta, Erdem Bangi

## Abstract

Cellular senescence is a conserved biological process essential for embryonic development, tissue remodeling, repair, and a key regulator of aging. Senescence also plays a crucial role in cancer, though this role can be tumor-suppressive or tumor-promoting, depending on the genetic context and the microenvironment. The highly heterogeneous, dynamic, and context-dependent nature of senescence-associated features and the relatively small numbers of senescent cells in tissues makes *in vivo* mechanistic studies of senescence challenging. As a result, which senescence-associated features are observed in which disease contexts and how they contribute to disease phenotypes remain largely unknown. Similarly, the specific mechanisms by which various senescence-inducing signals are integrated *in vivo* to induce senescence and why some cells become senescent while their immediate neighbors do not are unclear. Here, we identify a small number of cells that exhibit multiple features of senescence in a genetically complex model of intestinal transformation we recently established in the developing Drosophila larval hindgut epithelium. We demonstrate that these cells emerge in response to concurrent activation of AKT, JNK, and DNA damage response pathways within transformed tissue. Eliminating senescent cells, genetically or by treatment with senolytic compounds, reduces overgrowth and improves survival. We find that this tumor-promoting role is mediated by Drosophila macrophages recruited to the transformed tissue by senescent cells, which results in non-autonomous activation of JNK signaling within the transformed epithelium. These findings emphasize complex cell-cell interactions underlying epithelial transformation and identify senescent cell-macrophage interactions as a potential druggable node in cancer.

**One sentence summary:** Interactions between transformed senescent cells and macrophages drive tumorigenesis.

## INTRODUCTION

Cellular senescence is a highly conserved, dynamic, and regulated process crucial for proper embryonic development, tissue remodeling, repair, and aging (*1–5*). It is also a critical stress response mechanism induced by telomere shortening, DNA damage, oxidative stress, tissue damage, and aberrant activity of oncogenes(*5, 6*). Oncogene-induced senescence (OIS), triggered in response to hyperproliferation and transformation driven by cancer-driving genetic alterations, occurs independent of telomere erosion and is a crucial defense against cancer (*7*– *13*). This tumor suppressive role is primarily mediated by inducing a stable cell cycle arrest of transformed cells and promoting immune surveillance(*14, 15*). However, senescent cells can also promote tumorigenesis nonautonomously by remodeling the tumor microenvironment, which can result in increased proliferation, migration, and stemness of the neighboring non-senescent tumor cells and modulation of the immune system to make it more favorable for tumorigenesis(*14*).

Cellular senescence is broadly defined as a stable and generally irreversible growth arrest mediated by CDK inhibitors p21 and p16, a persistent DNA damage response, extensive heterochromatinization evident by the formation of Senescence Associated Heterochromatic Foci (SAHF), elevated senescence-associated β-gal activity (SA-β-gal) and a senescence-associated secretory phenotype (SASP) enriched in soluble factors with proliferative, immunomodulatory, proinflammatory and ECM remodeling activity(*1, 16*). Notably, features associated with senescence appear to be very heterogeneous, depending on the tissue, disease state, nature of the trigger, and time(*1, 17, 18*). In addition, not all senescent cells exhibit all features of senescence, and markers associated with senescence are not exclusive to this process(*19–21*). The highly heterogeneous nature of senescent cells, combined with their relatively low numbers in tissues, makes it particularly challenging to study how various stress signals are integrated at the tissue and cellular level to induce senescence *in vivo*(*22*). As therapeutic targeting of senescence will require the ability to clearly define and identify these cells in different disease contexts, the heterogeneity of the senescent cell phenotype also poses a significant barrier to biomarker discovery and drug development efforts(*17, 18*).

In addition to their heterogeneous features, the roles of senescent cells can vary widely depending on the disease and tissue contexts. For instance, senescence represents a powerful defense mechanism against cancer by inducing growth arrest of cells with activated oncogenes; a process demonstrated in experimental models and premalignant specimens from patients(*5, 23–25*). However, the persistence of senescent cells in malignant tissue can lead to strong pro-tumorigenic effects, including tumor relapse and increased malignancy(*26, 27*). The tumor-promoting effects of senescent cells are particularly alarming as many chemotherapies are known to induce senescence(*27*). The heterogeneous features and pleiotropic functions of senescence in cancer are further complicated by the genetically diverse nature of human tumors and complex interactions among tumor cells and their neighbors during tumorigenesis(*18*). Experimental models that can capture and explore this complexity are crucial for a comprehensive mechanistic understanding of context-dependent contributions of senescent cells to tumor progression.

We have previously leveraged similarities between the Drosophila and human intestine to study intestinal transformation by genetically manipulating Drosophila orthologs of genes recurrently mutated in colon tumors(*28, 29*). More recently, we established a 4-hit cancer model, KRAS TP53 PTEN APC, which expresses the oncogenic, mutant version of dRAS^G12V^ while simultaneously knocking down Drosophila orthologs of the recurrently mutated colorectal cancer tumor suppressors TP53, PTEN, and APC in the larval hindgut epithelium(*30*). APC, TP53, and KRAS are the three most frequently mutated genes in colon tumors(*31*). Similarly, the PI3K pathway, represented by PTEN loss in our model, is recurrently activated in many tumor types, including colon cancer, and is one of the most heavily targeted pathways in oncology drug development(*31, 32*), making this model representing a common colon cancer genome landscape a highly relevant experimental system to study senescence.

Multiple studies in Drosophila demonstrated the presence of senescent cells in various Drosophila tissues, indicating that cellular senescence is an evolutionarily conserved mechanism(*28, 33–36*). In this study, we show that cells that exhibit multiple features of senescence emerge in KRAS TP53 PTEN APC transformed hindguts, including nuclear accumulation of the Drosophila ortholog of p21, Dacapo (Dap), activation of DNA Damage Response (DDR) and formation of Senescence Associated Heterochromatic Foci (SAHF). We show that the senescent cell fate is determined in response to a multi-kinase network that includes the concurrent activation of AKT, JNK, and DDR signaling. Eliminating senescent cells from transformed tissue either genetically or by treatment with senolytic compounds(*37*) reduces tissue overgrowth and improve survival, demonstrating a tumor-promoting role for senescence in this genetic context. Our analysis shows that senescent cells recruit phagocytic hemocytes, the Drosophila macrophages(*38, 39*), to the transformed tissue by inducing MMP expression and compromising basement membrane integrity. Rather than clearing senescent cells and restoring tissue function, which is their normal function(*40*), macrophages recruited to the transformed tissue trigger a second phase of broader, non-autonomous JNK pathway activation in the transformed epithelium to further promote tumorigenesis.

## RESULTS

### Cells with multiple senescence markers emerge in KRAS TP53 PTEN APC hindguts

We have previously shown that targeting KRAS TP53 PTEN APC combination to the developing hindgut epithelium results in the expansion of the anterior region of the hindgut, where progenitor cells reside (Figure 1A,B) and increased proliferation (Figure 1C,D). Our analysis of KRAS TP53 PTEN APC transformed hindguts revealed a small number of cells expressing the Drosophila p21 ortholog Dacapo (Dap), which is undetectable in GFP-only control hindguts (Figure 1E). Additionally, we noted that subcellular localization of the Dap/p21 protein was different: some positive cells had nuclear localization of Dap/p21 while others had cytoplasmic (Figure 1F). p21 is a well-established marker of cellular senescence, suggesting that these cells may be senescent. A large body of prior work in multiple experimental systems has demonstrated that senescence is a dynamic and highly heterogeneous cellular state, emphasizing the importance of not relying on a single read-out to define cellular senescence. Therefore, we investigated whether Dap/p21 positive cells in KRAS TP53 PTEN APC hindguts exhibited other well-established senescence markers.

**Figure 1.**
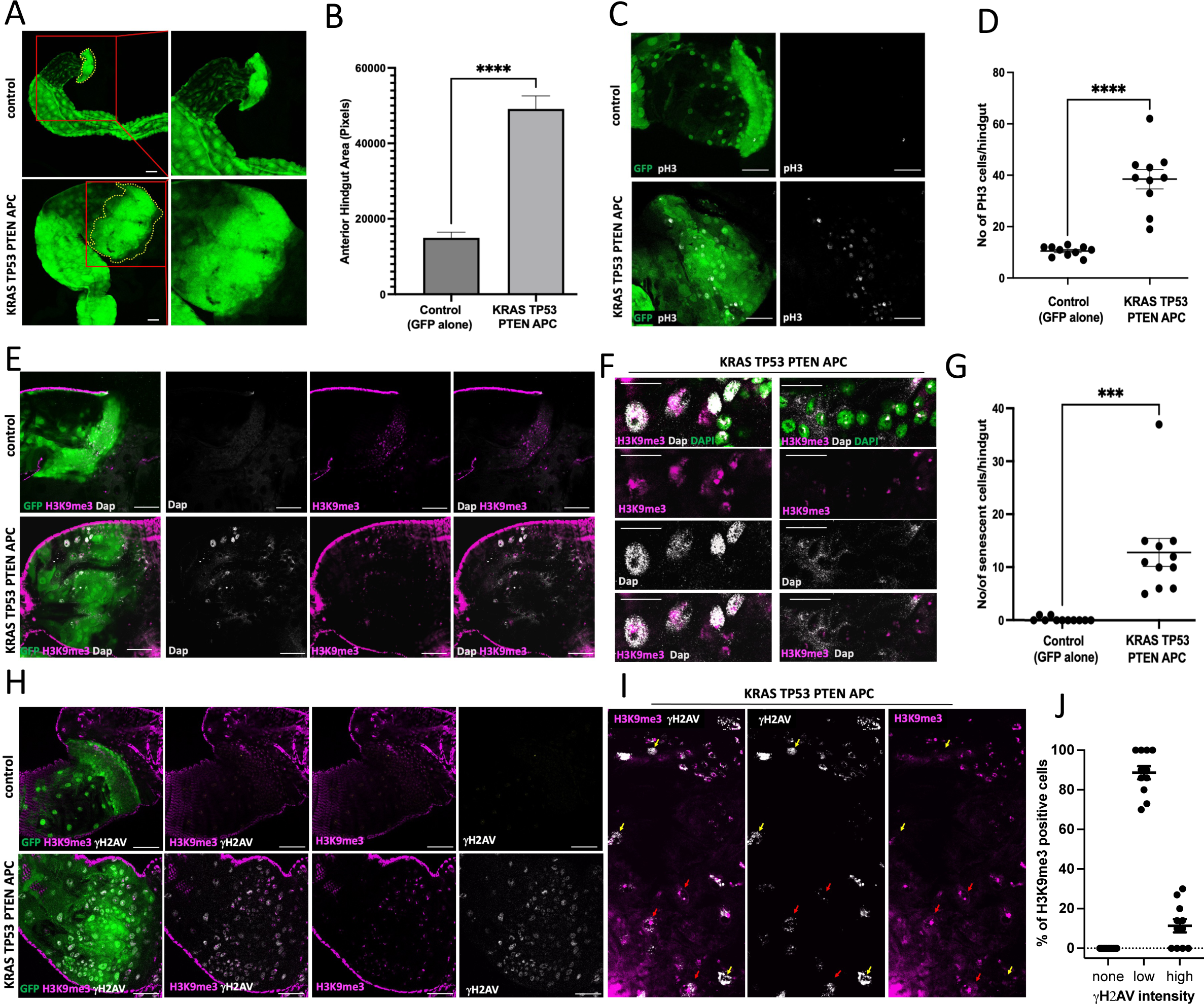
KRAS TP53 PTEN APC transformed hindguts display multiple markers of senescence. **(A)** Anterior hindgut area, marked with yellow dotted lines, in GFP-only control and KRAS TP53 PTEN APC larval hindguts. The right panels show magnified views of areas highlighted in red boxes in the left. **(B)** Quantification of larval anterior imaginal ring area of control versus KRAS TP53 PTEN APC transformed hindguts. **(C,D)** phospho-Histone 3 (pH3) staining (white) of the control and KRAS TP53 PTEN APC transformed hindgut epithelium and (green) **(C)** and quantification of the number of pH3+ cells/hindgut **(D)**. **(E)** Dap/p21 (white) and H3K9me3 (magenta) staining in hindguts (green) with indicated genotypes. **(F)** Magnified images of KRAS TP53 PTEN APC panels in **(E)** focusing on areas with nuclear and cytoplasmic Dap/p21 positive cells. **(G)** Quantification of the number of cells positive for nuclear Dap/p21 and H3K9me3 in hindguts with indicated genotypes. **(H)** Co-staining of H3K9me3 (magenta) and γ-H2AV (white) in hindgut epithelium (green) **(I)** Magnified areas of the KRAS TP53 PTEN APC panels in **(H)** Yellow and red arrows indicate cells with high and low γ-H2AV signal intensity, respectively. **(J)** Quantification of the fraction of H3K9me3 positive cells with low, high, or no γ-H2AV signal in KRAS TP32 PTEN APC hindguts. Error bars: Standard Error of the Mean (SEM). ***: p≤0.001, ****: p≤0.0001 (t-tests, PRISM Software). Scale bars: 50µM.

SAHF, another commonly observed feature of senescence, can be evaluated using the heterochromatin marker H3K9me3 (tri-methylated Histone 3 on Lysine 9)(*41*). KRAS TP53 PTEN APC transformed cells with nuclear Dap/p21 also had elevated levels of H3K9me3 visible throughout their nuclei; GFP-only controls or Dap/p21 negative cells in KRAS TP53 PTEN APC hindguts only exhibited the typical two foci that mark the mostly heterochromatic 4th chromosome pair in Drosophila cells (Figure 1E,F). Notably, while nuclear Dap/p21 and elevated H3K9me3 were fully concordant, none of the cells with cytoplasmic Dap/p21 exhibited elevated H3K9me3 (Figure 1F). We confirmed these results using RFP-HP1, another well-established marker of SAHF(*42, 43*)(Figure S1). Cytoplasmic localization of p21 has been observed in some tumors and may have an oncogenic function, but the mechanisms underlying this role remain unclear(*44, 45*). Whether cytoplasmic Dap/p21 plays a similar role in KRAS TP53 PTEN APC hindguts remains to be determined. Consistent with observations of senescent cells *in vivo*, the number of cells with nuclear Dap/p21 is low and variable, but they are present in all KRAS TP53 PTEN APC transformed hindguts (Figure 1G). We focused the rest of our analysis on these cells as they are also positive for two other well-established senescence markers.

Persistent DNA damage is another key feature of cellular senescence, commonly evaluated by the presence of γ-H2AX, which is produced by the phosphorylation of the histone variant H2AX and is a well-established read-out of DNA damage, repair, and senescence in mammalian systems and Drosophila. (*35, 46–48*). We found a high number of cells positive for γ-H2AV, the Drosophila equivalent of γ-H2AX, in KRAS TP53 PTEN APC transformed hindguts (Fig 1H). Colocalization studies using the SAHF marker H3K9me3 demonstrated that all H3K9me3 positive cells were also positive for γ-H2AV, with the majority exhibiting a low level of γ-H2AV signal intensity and a small number having high levels (Figure 1I,J). As all H3K9me3-high cells also have nuclear Dap/p21 (Figure 1E,F), this analysis demonstrates the presence of three well-established features of cellular senescence in these cells: Nuclear Dap/p21, formation of SAHF, and activation of DNA Damage Response (DDR). Significantly, not all γ-H2AV positive cells in KRAS TP53 PTEN APC transformed hindguts were positive for other senescence markers, consistent with previous studies that γ-H2AX is not an exclusive marker of cellular senescence. Our findings agree with previous work in mammalian models emphasizing the importance of using multiple markers to define senescent cells.

### Senescent cells promote KRAS TP53 PTEN APC-induced intestinal transformation

To determine whether senescence plays a role in KRAS TP53 PTEN APC-induced intestinal transformation, we sought to genetically eliminate senescent cells from the transformed tissue by knocking down Dap/p21. We found that reducing the level of Dap/p21 in KRAS TP53 PTEN APC hindguts was sufficient to prevent the formation of H3K9me3 positive foci, demonstrating that Dap/p21 is required for the emergence of SAHF, another key marker of senescence (Figure 2A). Dap/p21 knockdown in KRAS TP53 PTEN APC hindguts resulted in a significant reduction in the anterior hindgut expansion and rescue of lethality (Figure 2B,E, S2A). We found similar results upon feeding the p21 inhibitor UC2288(*49*) (Figure 2C,F, S2B). Furthermore, senolytic drugs navitoclax and fisetin, which have been shown to clear senescent cells in multiple experimental systems(*37*), also eliminated senescent cells in KRAS TP53 PTEN APC hindguts, reduced tissue expansion, and improved organismal survival (Figure 2D,G, S2C). Combined, these results demonstrate a tumor-promoting function for senescent cells in KRAS TP53 PTEN APC-induced intestinal transformation.

**Figure 2.**
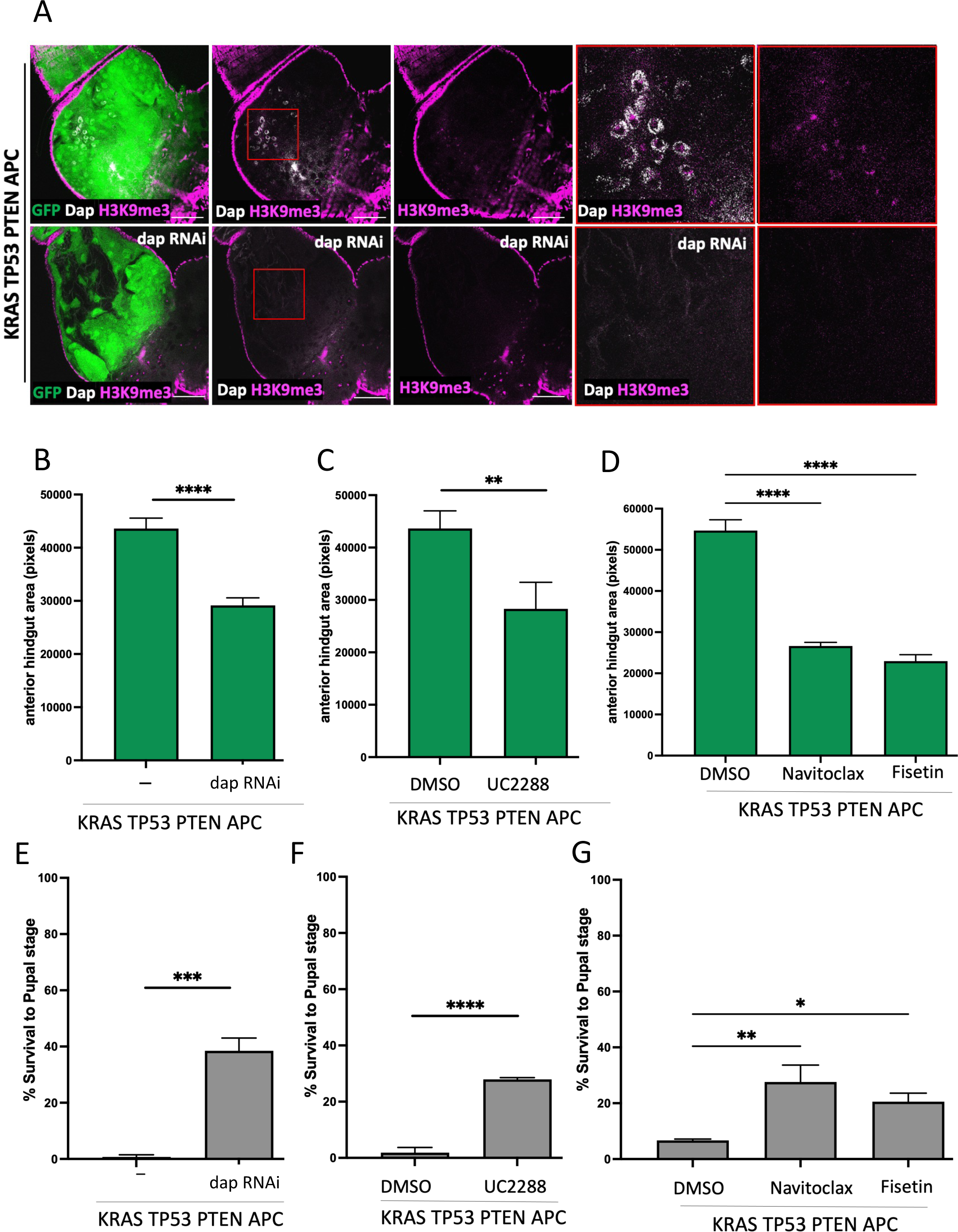
Senescent cells promote KRAS TP53 PTEN APC-induced intestinal transformation. **(A)** Dap/p21 (white) and H3K9me3 (magenta) staining of hindguts (green) with indicated genotypes. **(B-G)** The quantification of the anterior hindgut area **(B-D)** and survival to the pupal stage **(E-G)** of hindguts and experimental animals with indicated genotypes and treatment conditions. Error bars represent SEM. ****P<_0.0001, ***P<_0.001, **P<_0.01 (t-tests, PRISM Software) **(D,G)** Elimination of senescent cells by the senolytic compounds navitoclax and fisetin were sufficient to rescue reduction in anterior transformed hindgut area and KRAS TP53 PTEN APC induced larval lethality. Error bars: SEM. ****: p≤0.0001, **: p≤0.01, *: p≤0.05 (ANOVA, PRISM Software). Scale bars: 50µM.

### Senescence emerges in response to concurrent activation of AKT, JNK, and DDR signaling

Consistent with observations in human and other mammalian tissues, the number of Dap/p21 and H3K9me3 positive senescent cells in KRAS TP53 PTEN APC hindguts is low, and their location within transformed tissue is variable (Figure 1G), suggesting the presence of spatially restricted signals as determinants of the senescent phenotype. To understand how senescent cells emerge within transformed tissue and why some transformed cells become senescent while others in the same local microenvironment do not, we investigated whether signaling pathways activated in KRAS TP53 PTEN APC hindguts play a role as inducers of senescence.

We have previously demonstrated strong activation of AKT signaling in KRAS TP53 PTEN APC and other RAS/PI3K co-activated tumor models(*28, 30*) (Figure S3A). Reducing Akt levels in KRAS TP53 PTEN APC hindguts eliminated Dap/p21 expression (Figure 3A), demonstrating that AKT signaling has a conserved role in inducing Dap/p21 expression during transformation. However, AKT signaling is broadly activated throughout the transformed tissue (Figure S3A), making it unlikely to be the localized signal determining which transformed cells will become senescent.

**Figure 3.**
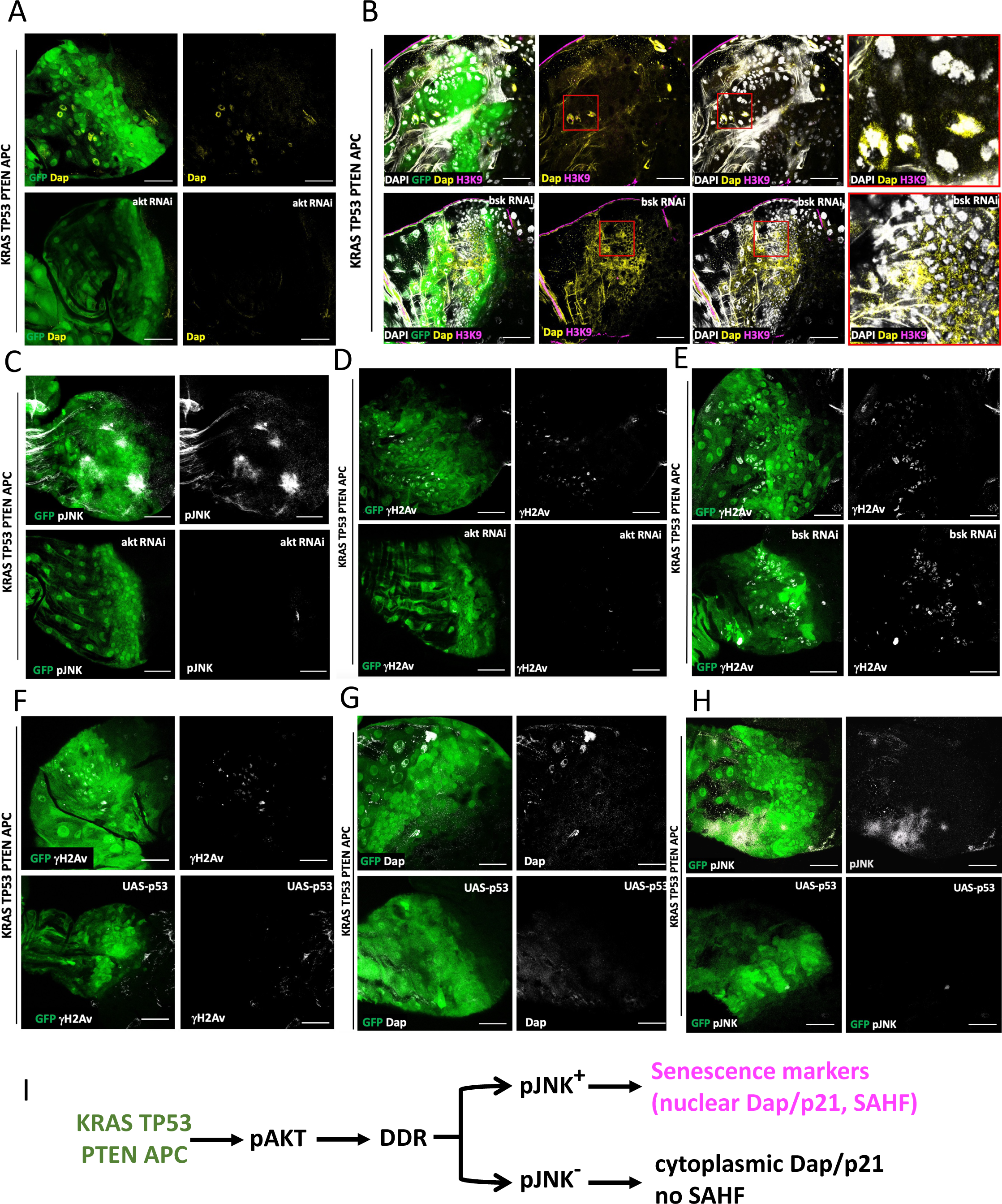
Determinants of senescence in KRAS TP53 PTEN APC transformed hindguts. **(A)** Dap/p21 staining (yellow) of KRAS TP53 PTEN APC transformed hindguts (green) with and without *akt* knockdown. **(B)** Dap/p21 (yellow) and H3K9me3 (magenta) co-staining of KRAS TP53 PTEN APC transformed hindguts (green) with and without *bsk* knockdown. No nuclear dap/p21 or H3K9me3 observed upon *bsk* knockdown. **(C,D)** phospho-JNK (pJNK, white, **C**) or γ-H2AV (white, **D**) staining of KRAS TP53 PTEN APC transformed hindguts (green) with and without *akt* knockdown. **(E)** γ-H2AV staining (white) of KRAS TP53 PTEN APC transformed hindguts (green) upon *bsk* knockdown. **(F-H)** γ-H2AV (white, **F**), Dap/p21 (white, **G**), and pJNK (white, **H**) staining of KRAS TP53 PTEN APC transformed hindguts (green) upon p53 overexpression. Scale bars: 50µM. **(I)** Working model: Senescence emerges in response to concurrent activation of AKT, JNK and DDR signaling (see main text for details).

Next, we explored a role for JNK signaling, which is activated in a more localized and variable manner in transformed tissue (Figure S3B). Knockdown of the Drosophila JNK ortholog *basket* (*bsk*) resulted in the complete elimination of both nuclear Dap/p21 and H3K9me3 as well as broad induction of cytoplasmic Dap/p21 (Figure 3B), suggesting that JNK signaling is required for Dap/p21 nuclear localization and senescence. As AKT signaling is required for both nuclear and cytoplasmic Dap/p21 (Figure 3A), we next tested whether AKT acts upstream of JNK signaling to induce Dap/p21 expression. Reducing AKT levels in KRAS TP53 PTEN APC hindguts resulted in a complete loss of JNK pathway activity (Figure 3E,F), demonstrating that JNK pathway activation is downstream of AKT signaling.

These findings demonstrate that only transformed cells with high AKT and JNK pathway activity become senescent. We hypothesize that JNK activity in pAKT-positive cells results in Dap/p21 nuclear localization and SAHF formation. However, the pattern of AKT activation is much broader than that of JNK signaling (Figure S3A,B), suggesting that not all pAKT-positive cells activate JNK signaling. Combined, these findings point to the involvement of another signal regulating Dap/p21 expression and nuclear localization.

Persistent DNA Damage Response (DDR) activation is a well-established feature of senescent cells, also conserved in our experimental system (Figure 1H-J). As DNA damage is a known activator of JNK signaling, we next investigated the relationship between AKT signaling, JNK signaling, and DDR. Reducing AKT levels in KRAS TP53 PTEN APC transformed hindguts resulted in a complete loss of γ-H2AV signal (Figure 3D), demonstrating that AKT activity is required to activate DDR. On the other hand, reducing JNK pathway activity by knocking down *bsk* did not affect γ-H2AV (Figure 3E), indicating that DDR is activated downstream of AKT signaling but does not require JNK signaling.

To determine whether persistent DDR signaling activates the JNK pathway in transformed tissue and induces senescence, we sought to genetically block DDR in KRAS TP53 PTEN APC hindguts. Overexpressing p53 in KRAS TP53 PTEN APC hindguts eliminated DDR-positive cells and induced apoptosis (Figure 3F, S3D). The loss of DDR activity in transformed tissue also prevented both Dap/p21 expression and JNK pathway activation. These findings demonstrate that the JNK pathway is activated downstream of DDR and that the senescent phenotype is an emergent feature of high pAKT, JNK, and DDR signaling (Figure 3I).

We hypothesize that AKT-dependent excessive proliferation in transformed hindgut tissue results in DNA damage in many transformed cells, broadly activating DDR. Persistent DDR results in the activation of JNK signaling in some of the cells, leading to nuclear Dap/p21 and other features of senescence. While all senescent cells exhibit DDR, not all DDR-positive cells become senescent. In addition, γ-H2AV signal intensity varies from cell to cell. The senescent phenotype largely correlates with a low level of γ-H2AV intensity, suggesting that JNK pathway activation and subsequent senescence may require a moderate threshold of DDR. These results demonstrate that senescence is a highly calibrated and integrated cellular response to excessive proliferation, DNA damage, oncogenic and stress signaling.

### A positive feedback loop between senescent cells and macrophages promotes intestinal transformation

Despite their low numbers within transformed tissue (Figure 1G), senescent cells appear to have a significant tumor-promoting effect (Figure 2). To investigate how senescent cells promote intestinal transformation, we tested whether they contribute to the expression of MMP1, which has been shown to be secreted by senescent cells in both mammalian systems and in Drosophila. KRAS TP53 PTEN APC transformed hindguts exhibit strong and broad MMP1 expression, which is eliminated upon Dap/p21 knockdown (Figure 4A), demonstrating that senescent cells are necessary for inducting MMP1 expression in KRAS TP53 PTEN APC transformed hindguts.

**Figure 4.**
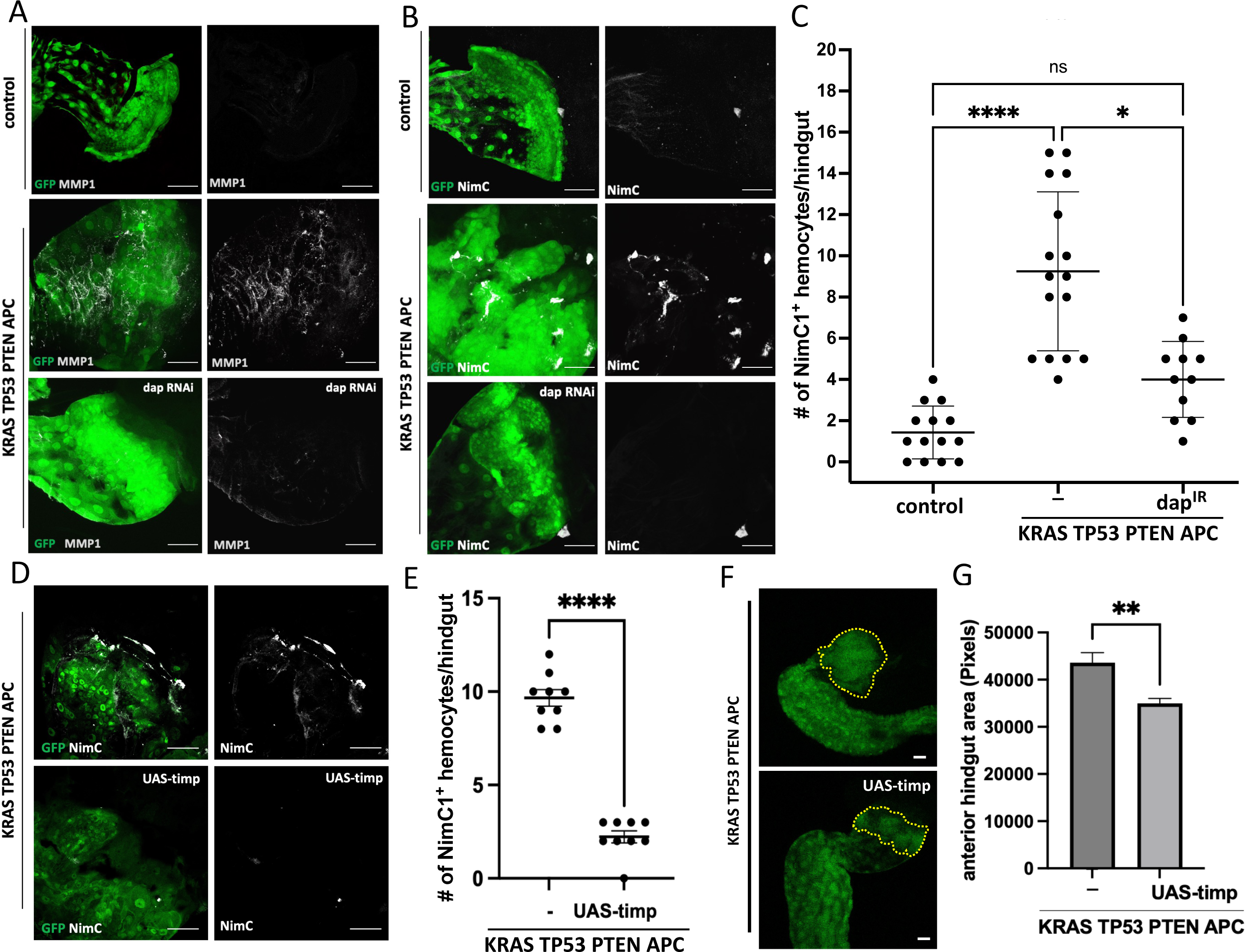
Mechanism of the tumor-promoting role of senescent cells. **(A,B)** MMP1 (white, **A**) and NimC1 (white, **B**) staining in hindguts (green) with indicated genotypes. **(C)** Quantification of the number of NimC1+ hemocytes associated with hindguts with indicated genotypes ****: p≤0.0001, *: p≤0.05 (ANOVA, PRISM Software). Error bars: SEM. **(D,E)** NimC1 staining (white, **C**) and quantification of the number of NimC1+ hemocytes associated with hindguts with indicated genotypes **(E)**. ****: p≤0.0001 (t-tests, PRISM Software). Error bars: SEM. **(F,G)** Anterior hindgut area, marked with yellow dotted lines in hindguts with indicated genotypes (green, **F**) and quantification of the anterior hindgut area size (**G**) p: **: p≤0.01 (t-tests, PRISM Software). Scale bars: 50µM.

A well-established function of senescent cells is to recruit macrophages to facilitate tissue remodeling and as a senescence surveillance mechanism to prevent the accumulation of senescent cells (*50–52*). As macrophages are often recruited to sites with compromised basement membrane integrity(*53, 54*), we tested whether senescent cells facilitate the recruitment of phagocytic hemocytes, the functional equivalent of mammalian macrophages (*38, 39, 55*), to the KRAS TP53 PTEN APC transformed tissue. We found that hemocytes, identified by the expression of the phagocytosis receptor NimC1 (*56, 57*), were recruited to KRAS TP53 PTEN APC transformed intestinal epithelium in a Dap/p21 dependent fashion (Figure 4B,C). Inhibition of MMP activity by ectopic expression of Timp, an endogenous inhibitor of the proteolytic activity of MMPs(*58*), significantly reduced the number of NimC1-positive hemocytes (Figure 4D,E). Blocking MMP activity and subsequent hemocyte recruitment also significantly reduced the size of the anterior hindgut area (Figure 4F,G). These findings suggest that senescent cells contribute to tumorigenesis by inducing MMP expression and hemocyte recruitment. Furthermore, disrupting basement membrane integrity by ectopically expressing MMP1 in an otherwise wildtype hindgut is sufficient to recruit hemocytes (Figure S4A,B), suggesting that senescence-induced MMP1 expression is the primary driver of hemocyte recruitment to the KRAS TP53 PTEN APC transformed tissue. Notably, hemocyte recruitment to an otherwise wildtype epithelium is not sufficient to induce transformation (Figure S4C,D), highlighting the context-dependent roles hemocytes play during tumorigenesis.

These findings demonstrate that senescent cells trigger MMP expression within KRAS TP53 PTEN APC transformed tissue, compromising basement membrane integrity and recruiting hemocytes. Given the broad MMP1 expression in KRAS TP53 PTEN APC transformed hindguts and the relatively low number of senescent cells, we next tested whether senescent cells induce MMP1 expression non-autonomously. As MMPs are direct transcriptional targets of JNK signaling(*59*), we reasoned that senescent cells may be expressing and secreting the Drosophila TNFα ortholog and the sole JNK pathway ligand Eiger(*60, 61*), to drive non-autonomous JNK pathway activation and MMP expression. Using the reporter construct *eiger-LacZ* (*62*), we found that *eiger* is not expressed in the transformed epithelium or the overlaying muscle; hemocytes were the only *eiger-LacZ* positive cells in KRAS TP53 PTEN APC transformed hindguts (Figure S5).

As senescent cells recruit hemocytes to the transformed tissue, any JNK signaling within the hindgut epithelium induced by hemocyte-derived Eiger would be downstream of senescence. However, we have previously shown that JNK signaling is also necessary to induce senescence (Figure 3B,I) and consequently hemocyte recruitment. These findings suggest a biphasic JNK pathway activation within the transformed epithelium, the first phase upstream of senescence, activated in a ligand-independent fashion in response to DDR, and required for the formation of senescent cells, MMP1 expression, and hemocyte recruitment. Eiger-expressing hemocytes then initiate the second phase of JNK pathway activation and MMP1 expression. To test this hypothesis, we knocked down the Eiger receptor Wengen (Wgn) in the hindgut epithelium, which would be expected only to affect the second, Eiger-induced phase of JNK signaling and MMP expression. Consistent with our hypothesis, *wgn* knock-down eliminated most of the MMP1 expression within KRAS TP53 PTEN APC transformed hindguts, only leaving a small domain that still exhibited a strong MMP signal (Figure 5A). We observed a similar pattern with JNK pathway activity, where after *wgn* knockdown, only a few pJNK-positive cells remained. Notably, the remaining MMP-positive (Figure 5A) and, pJNK-positive cells (Figure 5B) continued to exhibit strong signal intensity, indicating that these cells do not depend on hemocyte-derived Eiger for JNK pathway activation. In contrast, reducing the level of Bsk, the Drosophila c-Jun N-terminal Kinase (JNK), which is further downstream of the pathway and required for all JNK signaling, resulted in a much broader loss of pJNK throughout the hindgut (Figure S3B). We conclude that *wgn* knockdown eliminates the second phase, non-autonomous JNK pathway activation. In contrast, *bsk* knockdown removes both the DDR-induced first phase and the hemocyte-induced second phase of JNK signaling downstream of senescence. Consistent with this, we also observed a much stronger reduction in the anterior hindgut area upon *bsk* knockdown than *wgn* (Figure 5D,E).

**Figure 5.**
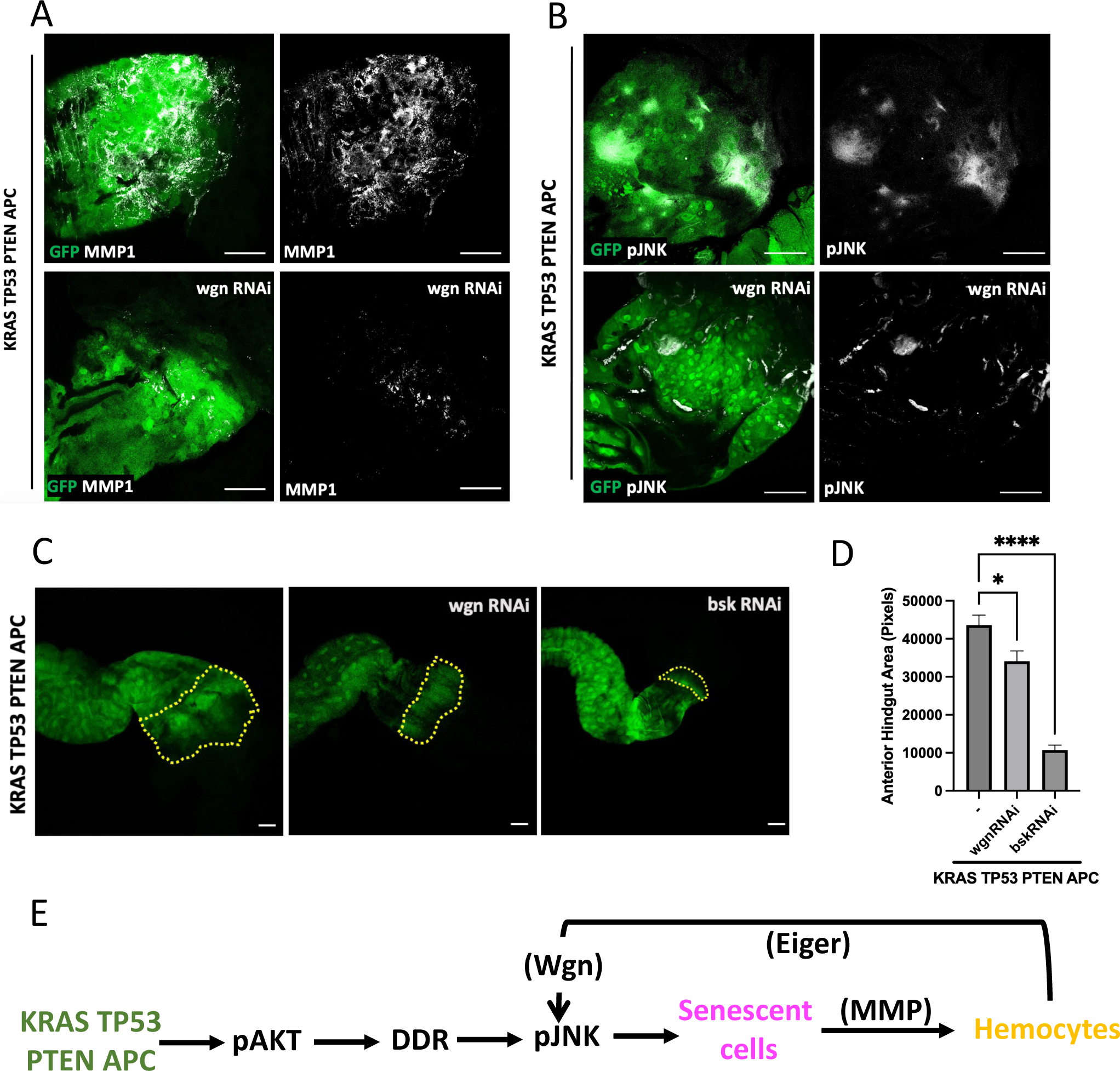
A positive feedback loop between senescent cells and hemocytes promotes intestinal transformation. **(A,B)** MMP1 (white, **A**) and pJNK (white, **B)** staining in hindguts with indicated genotypes. **(C)** Anterior hindgut area, marked with yellow dotted lines in hindguts with indicated genotypes (green, **C**) and quantification of the anterior hindgut area size **(D**). Error bars: SEM. **** p: ≤0.0001, *p: ≤0.05 (t-tests, PRISM Software). Scale bars: 50µM. **(E)** Working model: Reciprocal interactions between senescent cells in the transformed epithelium and recruited hemocytes recruited promotes tumorigenesis through biphasic activation of JNK signaling (see main text for details).

Overall, our findings highlight a positive feedback loop between senescent cells and hemocytes and biphasic JNK pathway activation as an important driver of tumorigenesis (Figure 5E). During intestinal transformation, concurrent activation of AKT, DDR, and JNK signaling triggers senescence in a small number of cells. Senescent-cell-driven MMP expression disrupts the basement membrane, resulting in the recruitment of phagocytic hemocytes, which express the JNK pathway ligand Eiger. Hemocyte-derived Eiger triggers a second phase of JNK pathway activation and MMP expression through its receptor Wgn in the transformed hindgut epithelium, further compromising basement membrane integrity and promoting hemocyte recruitment. Disrupting this feedback loop by eliminating senescent cells or blocking the Wgn-dependent second phase of JNK pathway activation suppresses tumorigenesis.

## DISCUSSION

Cellular senescence is a dynamic process regulated by complex interactions that integrate several short- and longer-range signals with intrinsic properties of individual cells, including their genetic makeup, disease state, or tissue of origin. Capturing and functionally exploring additional layers of complexity underlying the senescent cell fate and the cell non-autonomous effects of these cells during tissue homeostasis and disease development requires complex genetic manipulations. In this study, we leverage the powerful genetic tools available in Drosophila for a mechanistic exploration of triggers and consequences of senescence in a genetically complex model of colorectal cancer.

Multiple studies have shown that cells can respond to stress by undergoing cell death, cell repair, or senescence depending on the nature of the stress signal, cell type, and tissue microenvironment(*16, 63, 64*). The emergence of senescence depends on the dynamic interaction of various stressors acting on a tissue microenvironment. As most senescence triggers are typically broadly activated, elucidating how these signals cooperate *in vivo* to drive senescence in only a few cells has been difficult. Our study untangles the intricate interactions between multiple oncogenic and stress response pathways, including AKT signaling, JNK signaling, and the DDR pathway underlying the emergence of senescent cells in during intestinal transformation. We demonstrate that a cooperative epistatic interaction between these pathways leads to senescence and genetic targeting of any of these three pathways is sufficient to prevent the emergence of senescent cells.

Senescent tumor cells have been reported for multiple tumor types both in primary tumor specimens and in metastases (*14, 15*), though the specific triggers of senescence in different genetic contexts appear to be different and poorly understood. For instance, oncogenic KRAS-driven senescence is associated with DDR induced by reactive oxygen species(*65*). RAS/MAPK-PI3K coactivation, on the other hand, abrogates senescence and drives tumorigenesis in multiple experimental systems(*66, 67*). While p53 is a critical player in the induction of cell cycle arrest, senescence, and apoptosis in response to DNA damage and stress, p53-independent mechanisms of senescence have also been reported in mammalian cells(*68–70*). In this study, we demonstrate the emergence of senescent cells in KRAS TP53 PTEN APC transformed epithelium, which includes coactivation of RAS/MAPK and PI3K pathways and loss of p53 function, a commonly observed genetic landscape in multiple tumor types. It has been shown that AKT-induced senescence in p53 wildtype cells does not display DDR, SAHF, or high nuclear p16; instead, it depends on p53 (*66*). Notably, we find that AKT-dependent senescence in the absence of p53 function displays all these classical hallmarks of senescence. Our findings emphasize the importance of studying triggers and consequences of senescence in genetically complex and diverse cancer models to fully understand its nuanced role in tumor progression.

Recruitment of macrophages to sites of tissue damage by senescent cells is a well-documented senescence-surveillance mechanism to prevent the accumulation of senescent cells and promote tissue repair(*40, 71*). In multiple experimental systems, macrophage ablation leads to the accumulation of senescent cells, compromising tissue repair, remodeling, and embryonic development (*72–75*). Like senescent tumor cells, macrophages also play a crucial role in cancer, which can be tumor-promoting or tumor-suppressive depending on the tumor type, stage, genotype, and microenvironment (*76, 77*). However, the specific mechanisms mediating interactions between these cell types during tissue remodeling and cancer remain to be fully elucidated.

Tumor-associated hemocytes have also been reported in Drosophila cancer models, and their dual role in tumorigenesis is conserved (*78*). Furthermore, hemocyte-derived Eiger has been previously shown to both promote and suppress tumorigenesis in Drosophila models(*79–81*), but a potential role for senescence in these contexts has not been investigated. We demonstrate that senescent cells are the primary drivers of hemocyte recruitment by compromising basement integrity in an MMP-dependent manner. The TNFα ortholog Eiger secreted by these “tumor-associated hemocytes” is then co-opted to achieve more widespread JNK pathway activation and MMP expression within the transformed epithelium. These findings demonstrate how a comparatively low number of senescent cells in transformed tissue can non-autonomously trigger and maintain a self-sustaining positive feedback JNK signaling loop in cooperation with hemocytes to allow tumor growth and progression.

Targeting of senescent cells via senolytic compounds has been found to ameliorate disease states, including age-related disease as well as certain cancer types,(*82, 83*), although their mechanisms of action are still not well understood. Fisetin has been recently reported to reduce tumor progression in cancer patients and animal models (*84–86*); while navitoclax was reported to inhibit rapid tumor growth and development (*87*). We found that the senescent-cell clearing and anti-tumor effects of senolytic compounds are conserved in our experimental system, making it a powerful platform to explore the mechanisms of action of this promising class of cancer therapeutics. Furthermore, a better mechanistic understanding of interactions between senescent cells and macrophages offers opportunities to discover additional druggable nodes for therapeutic targeting of these cell-cell interactions in future studies.

## MATERIALS AND METHODS

### Fly Stocks

All Drosophila stocks were maintained on standard Drosophila media at room temperature. We have recently described the 4-hit model KRAS TP53 PTEN APC used in this study(*30*). *w^1118^* stock and UAS-based RNAi and overexpression lines were obtained from Bloomington Drosophila Stock Center (BDSC). The following stocks were used: *UAS-aktIR* (#31701, BDSC), *UAS-bskIR* (#31323, BDSC), *UAS-dapIR* (#36720, BDSC), *UAS-mmp1* (#58701, BDSC), *UAS-wgnIR* (#50594, BDSC), *UAS-timp* (#58708, BDSC), *RFP-HP1* (#30562, BDSC), *UAS-p53*(*88*) and *eiger-lacZ*(*62*) (gifted by K. Basler).

### Drosophila crosses to generate experimental and control animals

All crosses were set up on Bloomington’s semi-defined media. Experimental animals used in this study were generated and induced as previously described(*30*). Briefly, virgins were generated by heat-shocking *w^1118^ UAS-dcr2/Y, hs-hid; UAS-ras^G12V^ UAS-p53^RNAi^ UAS-pten^RNAi^ UAS-apc^RNAi^; byn-gal4 UAS-GFP tub-gal80^ts^ /S-T, Cy, tub-gal80, Hu, Tb* and *w^1118^ UAS-dcr2/Y, hs-hid; +; byn-gal4 UAS-GFP tub-gal80^ts^ /S-T, Cy, tub-gal80, Hu, Tb* stock to kill all male progeny as previously described(*30*). 18-20 virgins were then crossed to 10-12 males of either *w^1118^* as controls or those from stocks listed in the previous section. Crosses were kept at 18^0^C to keep transgenes silent during embryonic development and prevent early larval lethality. After 3 days of egg-laying, parents were removed and progeny were kept at 18^0^C for an additional 3 days to allow larval development. The transgenes were then induced by a temperature shift to 29^0^C for 3 days to generate the experimental and control animals used in this study.

### Dissections, immunofluorescence, and Microscopy

Hindguts from experimental and control larvae, identified by the absence of the Tubby (Tb) marker, were dissected and processed as previously described (*30*). Dissections were performed in 1X PBS, dissected tissue fixed with 4% paraformaldehyde, and processed for immunohistochemistry using our standard protocols(*28, 30*). The following primary antibodies were used: mouse anti-NimC1 (*79*)(Gifted by Eva Kurucz, 1:30), mouse anti-Mmp1(1:1000, DSHB #3B8D12), mouse anti-phospho JNK (1:50, Cell Signaling Technology #9255), mouse anti-phospho Akt (1:1000, Cell Signaling Technology #4054), mouse anti-Histone 2A gamma variant, phosphorylated (1:100, DSHB #UNC93-5.2.1), rabbit anti-Histone H3 (tri methyl K9) (1:50, Abcam #ab8898), mouse anti-Dacapo (1:10, DSHB #NP1), rabbit anti-Dcp1(1:1000, Cell Signaling Technology #9578), mouse anti-β-gal (1:100, DSHB #40-1a), rabbit anti-β-gal (1:200, Thermo Fisher Scientific #A111-32). Alexa-conjugated goat-anti-mouse and anti-rabbit antibodies were used as secondary antibodies (1:1000, Thermo Fisher Scientific, #A-11031, #A-21052, #A-110356, #A-21071). All guts were imaged using SPE DM6 Leica Confocal Microscope at 40X magnification at 1.0 Zoom. Hindguts used to quantify the imaginal ring proliferation area were imaged at 10X magnification, 1.5X Zoom (*30*).

### Scoring and Quantification

Crosses for survival studies were set up in triplicates with 18-20 virgin females of KRAS TP53 PTEN APC crossed to 10-12 males at 29^0^C and parents were removed after 2 days of egg laying. KRAS TP53 PTEN APC virgins crossed to *w^1118^* males were used as baseline controls. The progeny were allowed to develop for the next 10 days at 29^0^C and survival to pupal stage was calculated by counting the number of experimental and control pupae as previously described(*30*). The size of the anterior hindgut area was measured using Image J(*30*). Cell number quantifications were performed on Leica SPE DM6 confocal microscope at 40X magnification. Statistical analysis was done using PRISM software.

### Drug Administration

Drug concentrations used in this study are as follows: navitoclax (10uM Selleck Chemicals #S1001), fisetin (10uM, Selleck Chemicals #S2298), UC2288 (10uM, Millipore Sigma/ Sigma-Aldrich #532813). Drugs were diluted in Bloomington semi-defined media to achieve the indicated final concentrations in the food along with 0.1% dimethyl sulfoxide (DMSO) as previously described(*29*). Food with 0.1% DMSO alone was used as a vehicle-only control.

### Supplemental Materials

**Supplementary Figure 1.**
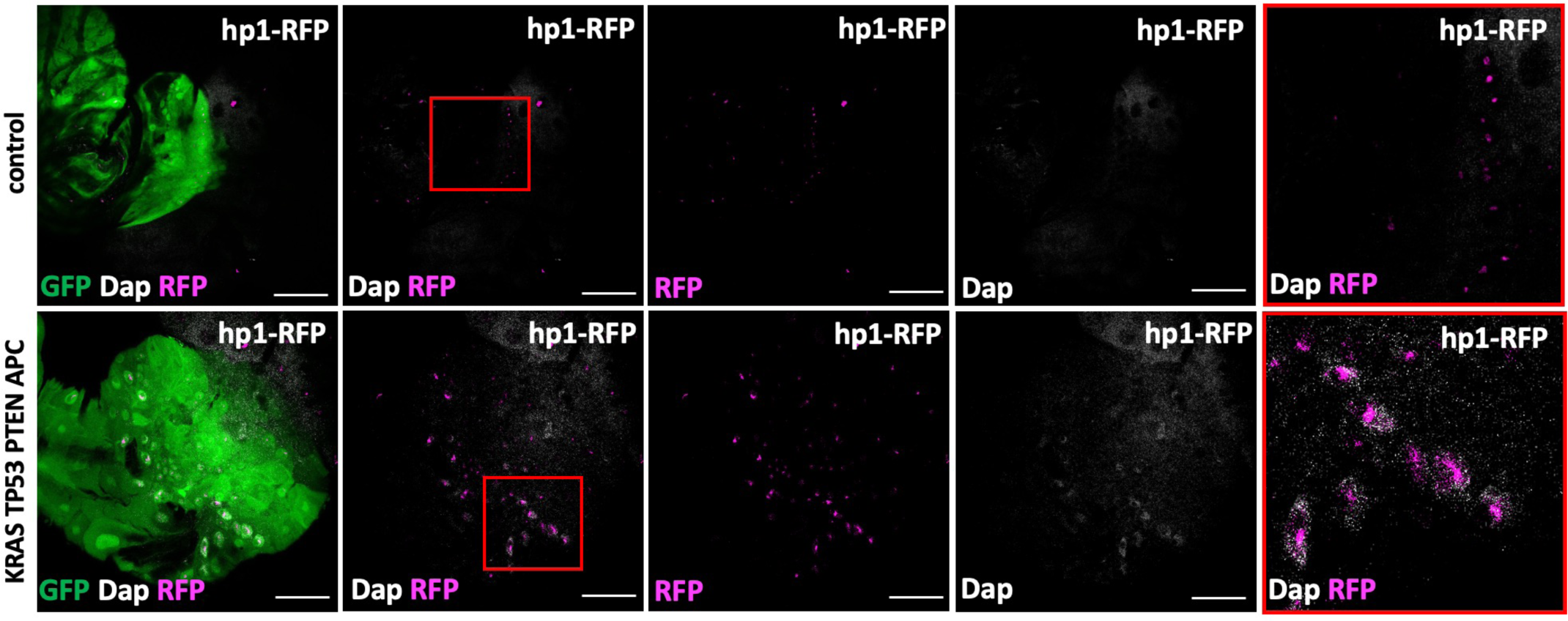
KRAS TP53 PTEN APC transformed hindguts display another marker of SAHF. Colocalization of nuclear Dap/p21 (white) and RFP-HP1 (magenta), another marker of heterochromatin and SAHF in KRAS TP53 PTEN APC transformed hindguts. GFP only control hindgut epithelium does not show Dap/p21 or RFP-HP1 positive cells Scale bars: 50µM.

**Supplementary Figure 2.**
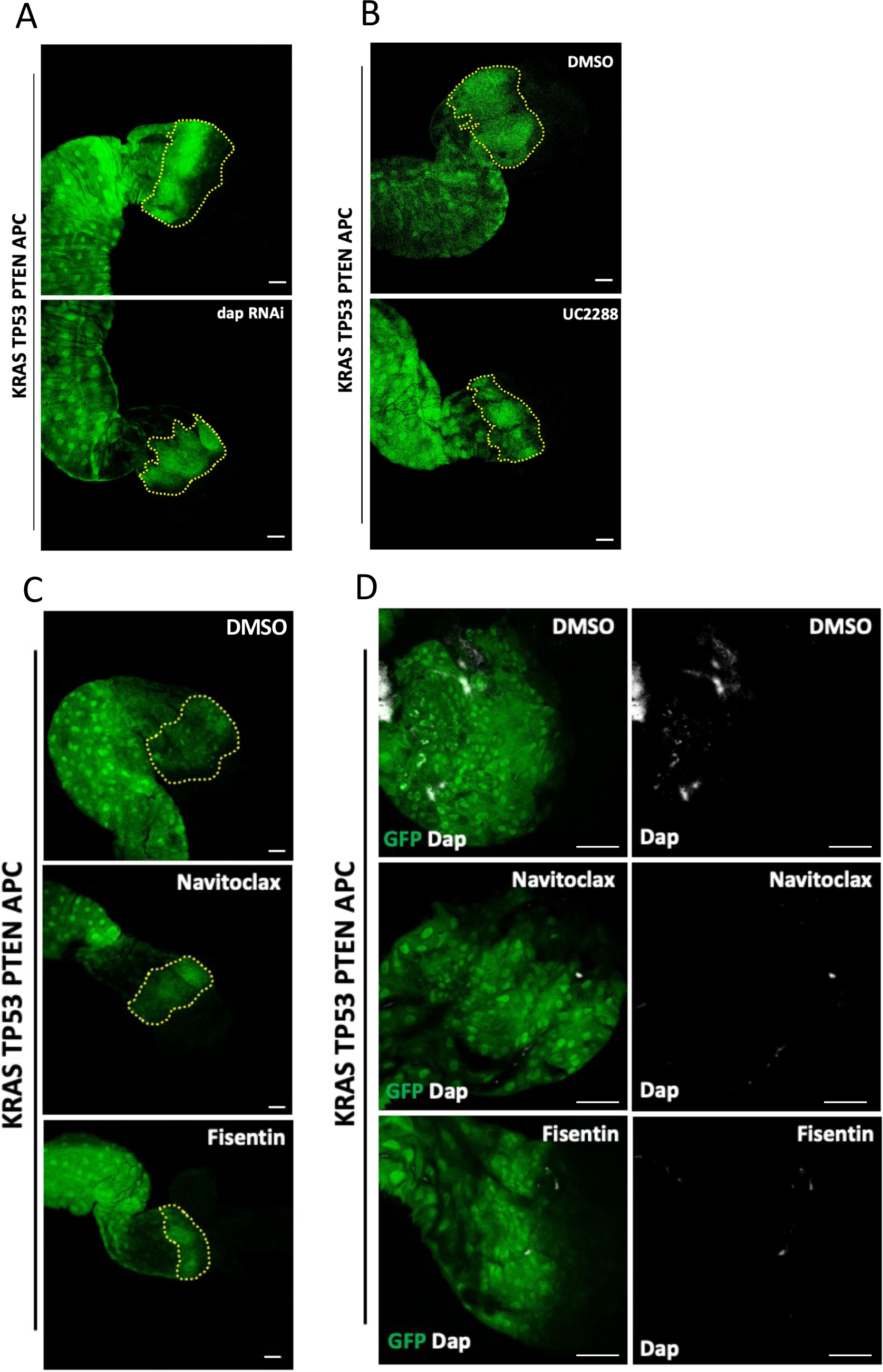
Representative hindgut images used for quantifications in Figure 2. **(A-C)** Example KRAS TP53 PTEN APC hindguts (green) with and without Dap/p21 knockdown, treated with DMSO only and UC2288 **(B)**, treated with DMSO only, Navitoclax and Fisetin **(C)**. The yellow dotted lines in **(A-C)** indicate the larval imaginal ring area quantified for the analysis presented in Figure 2. **(D)** Dap/p21 staining (white) in KRAS TP53 PTEN APC (green) hindguts treated with DMSO, navitoclax and fisetin. Scale bars : 50µM.

**Supplementary Figure 3.**
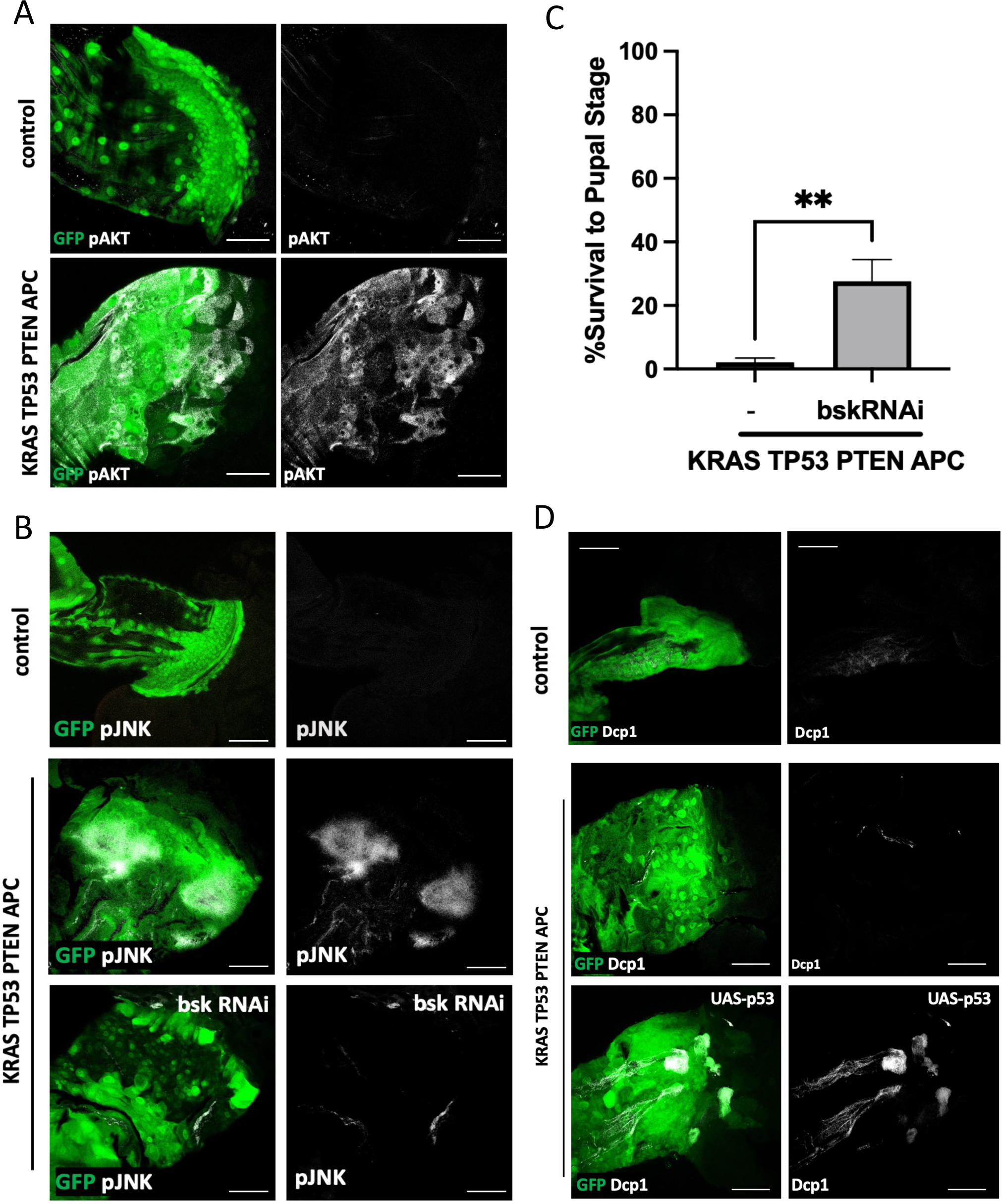
Evaluating the activity of senescence-relevant pathways in KRAS TP53 PTEN APC transformed hindguts. **(A)** phospho-AKT (pAKT, white) staining in GFP only control and KRAS TP53 PTEN APC hindgut epithelium (green). **(B)** phospho-JNK (pJNK, white) staining in hindguts (green) with indicated genotypes. **(C)** Quantification of survival to pupal stage of of animals with indicated genotypes Rescue of KRAS TP53 PTEN APC induced larval lethality with bsk RNAi. Error bars : SEM. **: p≤0.01, (t-tests, PRISM Software) **(D)** Cleaved effector caspase Dcp1 staining (white), a well-established read-out of apoptosis, in hindguts (green) with indicated genotypes. Scale bars: 50µM.

**Supplementary Figure 4.**
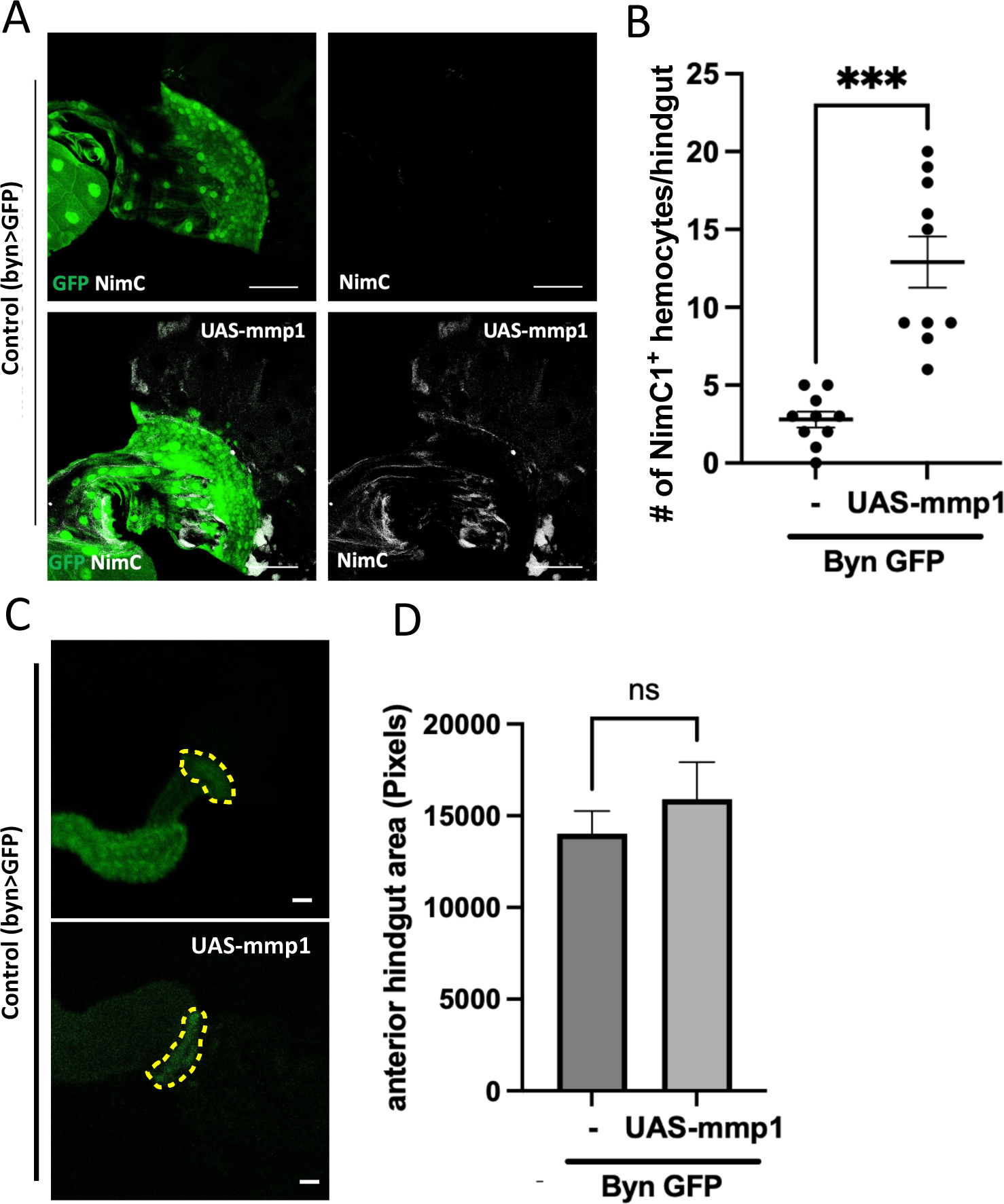
MMP1 expression in wildtype hindgut epithelium is sufficient to drive hemocyte recruitment. **(A,B)** NimC1 staining (white) in GFP-only or MMP1 expressing hindguts and the quantification of the number of NimC1+ hemocytes associated with each hindgut Error bars: SEM. ***p: ≤0.001 (t-tests, PRISM software). **(C)** Anterior hindgut area, marked with yellow dotted lines in hindguts with indicated genotypes (green, **C**) and quantification of the anterior hindgut area size **(D).** Error bars represent SEM. (t-tests, PRISM software). Scale bars: 50µM.

**Supplementary Figure 5.**
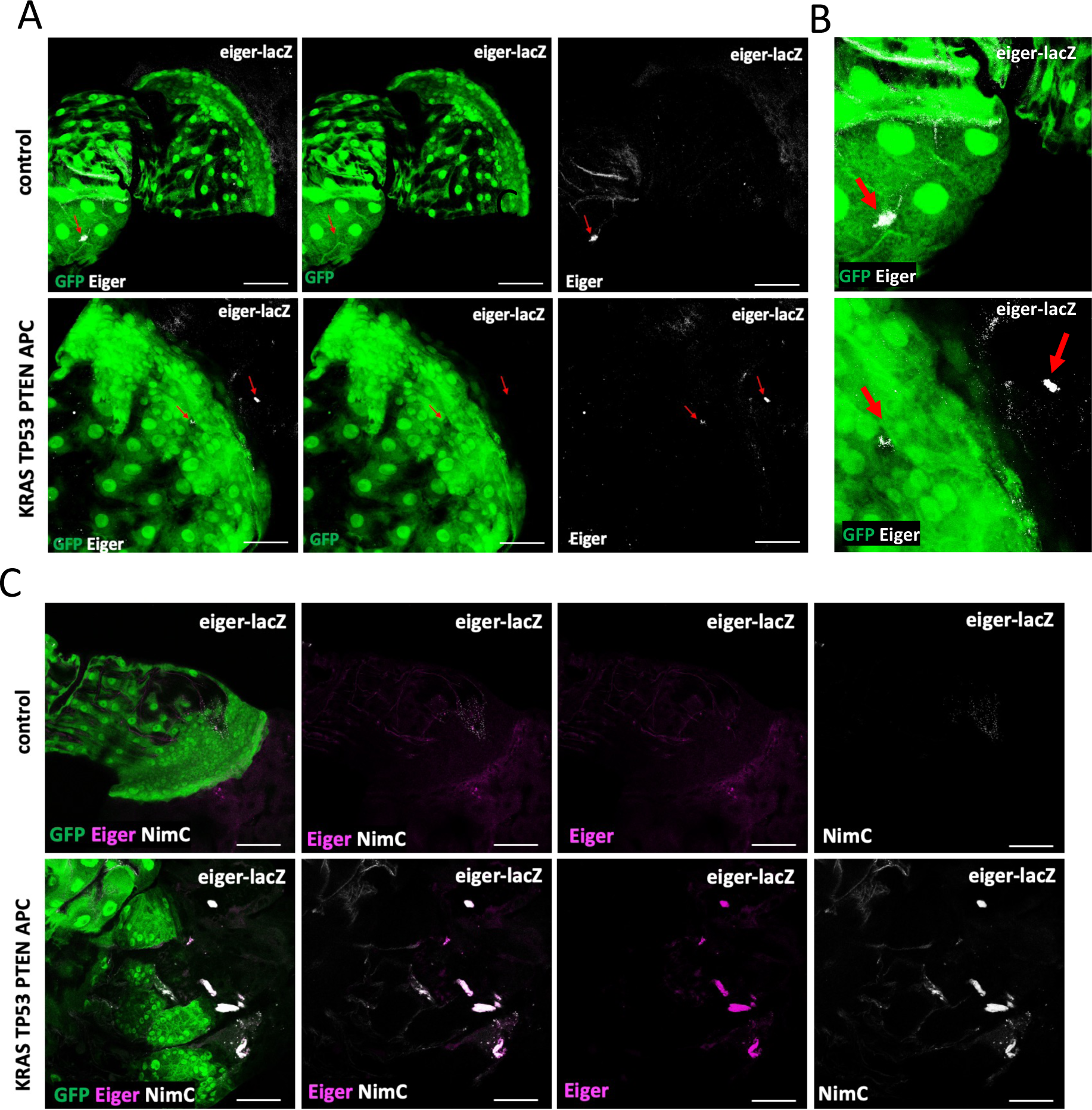
Hemocyte recruitment to the KRAS TP53 PTEN APC transformed hindgut epithelium. **(A)** β-gal staining (to detect eiger-LacZ, white) in hindguts with indicated genotypes. **(B)** Enlarged images of the panels in **(A)** showing that *eiger-LacZ* (white) is not expressed in the control or transformed hindgut epithelium (green). Red arrows: *eiger-LacZ* positive cells. **(C)** Co-staining for *eiger-LacZ* (magenta) and NimC1 (white) in KRAS TP53 PTEN APC in hindguts with indicated genotypes (green) showing co-localization for *eiger-LacZ* and NimC1. Scale bar : 50µM.

## Acknowledgments

This study used monoclonal antibodies obtained from the Developmental Studies Hybridoma Bank, created by the NICHD of the NIH and maintained at The University of Iowa, Department of Biology, Iowa City, IA 52242, transgenic Drosophila RNAi lines (Office of the Director R24 OD030002: “TRiP resources for modeling human disease”) and other lines obtained from the Bloomington Drosophila Stock Center (NIH P40OD018537) and the Vienna Drosophila Resource Center (VDRC, www.vdrc.at). We thank Juan Martin Portilla, Autumn Hawkins and Tajah Vassel for technical support, Dr. Konrad Basler and Dr. Eva Kurucz for sharing reagents.

## Author contributions

Supervision, funding acquisition, project administration: EB. Conceptualization, methodology, writing, reviewing, and editing : EB, ID. Investigation, visualization: ID

## Data and materials availability

All data are available in the main text or the supplementary materials.

